# Comprehensive bioinformatics analysis of *L1CAM* gene revealed Novel Pathological mutations associated with L1 syndrome

**DOI:** 10.1101/561431

**Authors:** Naseem S. Murshed, Mujahed I. Mustafa, Abdelrahman H. Abdelmoneim, Thwayba A. Mahmoud, Nafisa M. Elfadol, Mohamed A. Hassan

**Affiliations:** Department of Biotechnology, Africa city of Technology, Sudan; Department of Biochemistry, University of Bahri, Sudan

**Keywords:** L1CAM, L1 syndrome (CRASH syndrome), Comprehensive bioinformatics analysis, SNPs, diagnostic markers, breast cancer

## Abstract

**Background:** Mutations in the human L1CAM gene cause a group of neurodevelopmental disorders known as L1 syndrome (CRASH syndrome). The L1CAM gene provides instructions for producing the L1 protein, which is found all over the nervous system on the surface of neurons. L1 syndrome involves a variety of characteristics but the most common characteristic is muscle stiffness. Patients with L1 syndrome can also suffer from difficulty speaking, seizures, and underdeveloped or absent tissue connecting the left and right halves of the brain.

**Method:** The human L1CAM gene was studied from dbSNP/NCBI, 1499 SNPs were Homo sapiens; of which 450 were missense mutations. This selected for Comprehensive bioinformatics analysis by several in silico tools to investigate the effect of SNPs on L1CAM protein’s structure and function.

**Results:** 34 missense mutations (26 novel mutations) out of 450 nsSNPs that are found to be the most deleterious that effect on the L1CAM structural and functional level.

**Conclusion:** Better understanding of L1 syndrome caused by mutations in L1CAM gene was achieved using Comprehensive bioinformatics analysis. These findings describe 35 novel L1 mutations which improve our understanding on genotype-phenotype correlation. And can be used as diagnostic markers for L1 syndrome and besides in cancer diagnosis specifically in breast cancer.

## 1. Introduction

L1 syndrome (also known as CRASH syndrome) is a group of a very rare inherited disorders that primarily affect the nervous system and is characterized by Hydrocephalus with Stenosis of the Aqueduct of Sylvius (HSAS), intellectual disability, corpus callosum hypoplasia (or agenesis), adducted thumbs and spastic paraplegia (1-4). It is a recessive X-linked disorder that is exclusively affects men with incidence of 1/30.000 male births (5), and it is caused by mutations in the *L1CAM* gene (6-19) located near the telomere of the long arm of X chromosome in Xq28 in humans (20-22). Over 200 mutations in L1CAM gene have been reported (3, 17), this gene encodes for the L1 Cell Adhesion Molecule protein, which is a member of the immunoglobulin superfamily. As the name implies; the protein enables the adhesion of neural cells to one another and it is a key regulator of synapse formation, synaptic plasticity and axons and dendrites growth and formation (23-26). It was found that *L1CAM* gene is a major driver for tumor cell invasion and motility (27) therefore fully understanding its function will aid in the diagnosis and treatment of different types of cancers (27-50).

The underline pathogenetic mechanisms by which L1 syndrome happens remains unsolved (6, 25, 51), and the treatment requires shunting of the cerebrospinal fluid as needed (52). We hope for a better understanding of the condition and we believe thorough investigation of the *L1CAM* gene might help in that.

The aim of this study is to identify pathogenic mutations in the coding region of *L1CAM* gene using variant bioinformatics tools, which might then be used as diagnostic markers and may help in the development of new therapeutic strategies using gene therapy and pharmacogenomics. This is the first in silico analysis in the coding region of *L1CAM* gene that prioritized the functional analysis of nsSNPs. The use of variant bioinformatics tools was extremely beneficial due to the elimination of the cost and conformation of the results by the different-parameters-based softwares and It will facilitate the future genetic studies (53).

## 2. Materials and Methods

### 2.1 Data mining

The data on human *L1CAM* gene was collected from National Center for Biological Information (NCBI) web site (54). (https://www.ncbi.nlm.nih.gov/) and the protein sequence was collected from Uniprot (55) (https://www.uniprot.org/).

### 2.2 SIFT

We used SIFT to observe the effect of A.A. substitution on protein function. SIFT predicts damaging SNPs on the basis of the degree of conserved amino acid residues in aligned sequences to the closely related sequences, gathered through PSI-BLAST (56, 57). It’s available at (http://sift.jcvi.org/).

### 2.3 PolyPhen

We used PolyPhen (version 2) to study the probable impacts of A.A. substitution on structural and functional properties of the protein by considering physical and comparative approaches (58, 59). It is available at (http://genetics.bwh.harvard.edu/pph2/).

### 2.4 PROVEAN

PROVEAN is a software tool which predicts whether an amino acid substitution or indel has an impact on the biological function of a protein. It is useful for filtering sequence variants to identify nonsynonymous or indel variants that are predicted to be functionally important (60, 61). It is available at (https://rostlab.org/services/snap2web).

### 2.5 SNAP2

SNAP2 is a trained classifier that is based on a machine learning device called “neural network”. It distinguishes between disease-associated and neutral variants/non-synonymous SNPs by taking a variety of sequence and variant features into account. It is available at (https://rostlab.org/services/snap2web/).

### 2.6 SNPs&GO

SNPs&GO is an accurate method that, starting from a protein sequence, can predict whether a variation is disease related or not by exploiting the corresponding protein functional annotation. (62, 63). It is available at (http://snps.biofold.org/snps-and-go/snps-and-go.html).

### 2.7 PHD-SNP

An online Support Vector Machine (SVM) based classifier, is optimized to predict if a given single point protein mutation can be classified as disease-related or as a neutral polymorphism. It is available at: (http://snps.biofold.org/phd-snp/phdsnp.html).

### 2.8 I-Mutant 3.0

I-Mutant 3.0 Is a neural network based tool for the routine analysis of protein stability and alterations by taking into account the single-site mutations. The FASTA sequence of protein retrieved from UniProt is used as an input to predict the mutational effect on protein stability (64). It is available at (http://gpcr2.biocomp.unibo.it/cgi/predictors/I-Mutant3.0/I-Mutant3.0.cgi).

### 2.9 MUpro

MUpro is a support vector machine-based tool for the prediction of protein stability changes upon nsSNPs. The value of the energy change is predicted, and a confidence score between −1 and 1 for measuring the confidence of the prediction is calculated. A score <0 means the variant decreases the protein stability; conversely, a score >0 means the variant increases the protein stability. It’s available at (http://mupro.proteomics.ics.uci.edu/).

### 2.10 GeneMANIA

We submitted genes and selected from a list of data sets that they wish to query. GeneMANI approach to know protein function prediction integrate multiple genomics and proteomics data sources to make inferences about the function of unknown proteins (65-67). It is available at (http://www.genemania.org/).

### 2.11 Identification of Functional SNPs in Conserved Regions by using ConSurf server

ConSurf web server provides evolutionary conservation profiles for proteins of known structure in the PDB. Amino acid sequences similar to each sequence in the PDB were collected and aligned using CSI-BLAST and MAFFT, respectively. The evolutionary conservation of each amino acid position in the alignment was calculated using the Rate 4Site algorithm, implemented in the ConSurf web server. It is available at (http://consurf.tau.ac.il/).

### 2.12. Structural Analysis

#### 2.12.1 Detection of nsSNPs Location in Protein Structure

Mutation3D is a functional prediction and visualization tool for studying the spatial arrangement of amino acid substitutions on protein models and structures. Further, it presents a systematic analysis of whole genome and whole-exome cancer datasets to demonstrate that mutation3D identifies many known cancer genes as well as previously underexplored target genes (68). It is available at (http://mutation3d.org).

### 2.13 Analysis of 3 UTR and 5 UTR of L1CAM gene

Sequence related to the 3 UTR and 5 UTR of *L1CAM* gene were retrieved from ensemble website, (https://www.ensembl.org/index.html). which were inserted in to RegRNA 2 website to generate the related microRNA sequences. (http://regrna2.mbc.nctu.edu.tw/) These results are also given to miRmap software to get free energy and conservation values. (https://mirmap.ezlab.org/).

## 3. Results

## 4. Discussion

26 novel mutations were found to have a damaging effect on the stability and function of the *L1CAM* gene using Comprehensive bioinformatics analysis tools. L1 Cell Adhesion Molecule (*L1CAM*) gene is a Protein Coding gene that plays a role In the axon outgrowth and path finding during the development of the nervous system. It is specialized extensions of neurons that transmit nerve impulses. *L1CAM* has been identified in a group of overlapping X-linked neurological disorders known as L1 syndrome (52).

Many *Homo sapiens* SNPs that are now recognized (https://www.ncbi.nlm.nih.gov/snp), open the doors to improve our understanding on genotype-phenotype correlation. Therefore, a deep comprehensive bioinformatics analysis was made to prioritize SNPs that have a structural and functional impact on L1CAMprotein. The most frequent type of genetic mutation is the single nucleotide polymorphism (SNP). Non-synonymous SNPs (nsSNPs) or missense mutations arise in coding region. nsSNPs result in a single amino acid substitution which may have effects on the structure and/or function of protein (69). Therefore, in this study we focus on SNPs in coding and non-coding regions. We investigated the effect of each SNP on the function and stability of the protein and gene expression of related genes using different bioinformatics tools with different parameters and aspects, in order to confirm the results we found and to minimize the error to the least percentage possible. The software used were SIFT, Polyphen-2, PROVEAN, SNAP2, SNP&GO, PhD-SNP, I-Mutant 3.0, MUPro and Mutation3D (Figure 1)

**Figure 1:**
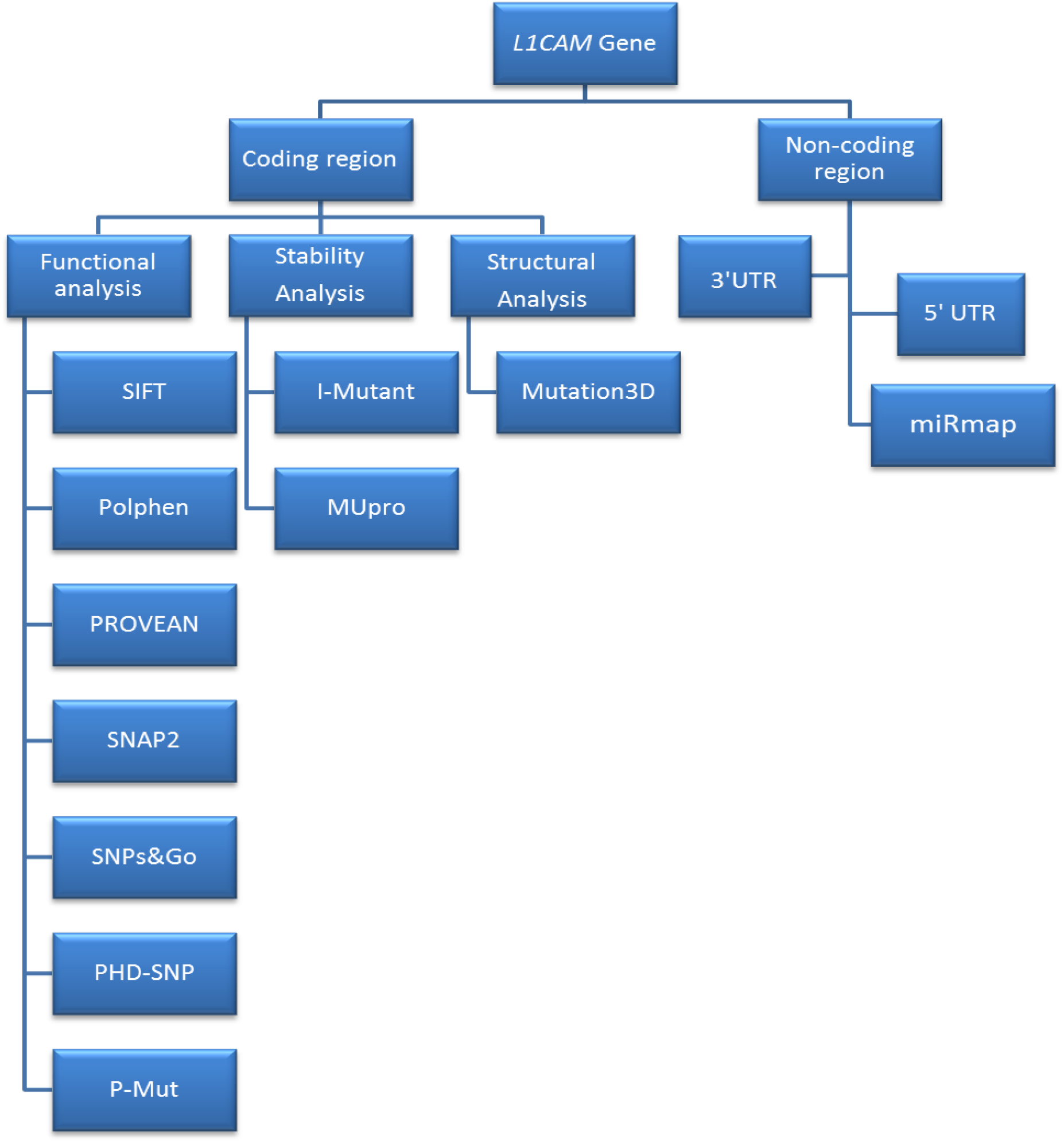
Diagrammatic representation for *L1CAM* gene in coding region in silico work flow.

1498 SNPs were retrieved from the dbSNP/NCBI Database, which was the total number of nsSNPs in the coding region of the *L1CAM* gene. There were 450 nsSNPs (missense mutations) then submitted them to functional analysis by SIFT, PolyPhen-2, PROVEAN and SNAP2. SIFT server predicted 140 deleterious SNPs, polyphen-2 predicted 224 damaging SNPs (75 possibly damaging and 149 probably damaging), PROVEAN predicted 146 deleterious SNPs while in SNAP2 we filtered the triple-positive deleterious SNPs from the previous three analysis tools, out of 49, There were 43 nsSNPs predicted deleterious SNPs by SNAP2. (Table 1) After filtering the Quadra-positive deleterious SNPs we ended up with 43 SNPs and we submitted them to SNPs&GO and PhD-SNP to further investigate their effect on the functional level. PhD-SNP predicted 37 disease-associated SNPs while SNP&GO predicted 38, so we filtered the double positive 34 SNPs (Table 2) and submitted them to I-Mutant 3.0, P-MUT and MUPro respectively (Table 3) to investigate their effect on structural level. All the SNPs were found to cause a decrease in the stability of the protein except for six SNPs predicted by I-Mutant 3.0 to increase the stability, one SNP (P240L) prediction by MUPro and the SNPs in P-MUT predicted 34 deleterious.

**Table1:** Damaging (Deleterious) nsSNPs associated variations predicted by various softwares:

| SUB | SIFT<br>prediction | Score | PolyPhen<br>prediction | Score | PROVEAN<br>prediction | Score | SNAP2<br>prediction | Score |
| --- | --- | --- | --- | --- | --- | --- | --- | --- |
| E1175K | Affect | 0 | probably damaging | 1 | Deleterious | -3.609 | Effect | 63 |
| K1150R | Affect | 0 | probably damaging | 1 | Deleterious | -2.81 | Effect | 42 |
| G1149R | Affect | 0 | probably damaging | 1 | Deleterious | -7.494 | Effect | 59 |
| R1145H | Affect | 0 | probably damaging | 1 | Deleterious | -4.506 | Effect | 76 |
| R1145C | Affect | 0 | probably damaging | 1 | Deleterious | -7.228 | Effect | 66 |
| L1135V | Affect | 0 | probably damaging | 1 | Deleterious | -2.844 | Effect | 38 |
| L1132P | Affect | 0 | probably damaging | 1 | Deleterious | -6.585 | Effect | 13 |
| R990C | Affect | 0 | probably damaging | 1 | Deleterious | -4.943 | Effect | 41 |
| N945I | Affect | 0 | probably damaging | 1 | Deleterious | -8.052 | Effect | 44 |
| N825K | Affect | 0 | probably damaging | 1 | Deleterious | -5.312 | Effect | 46 |
| N825Y | Affect | 0 | probably damaging | 1 | Deleterious | -7.206 | Effect | 48 |
| P816S | Affect | 0 | probably damaging | 1 | Deleterious | -7.806 | Effect | 43 |
| Y784C | Affect | 0 | probably damaging | 1 | Deleterious | -8.014 | Effect | 65 |
| F781L | Affect | 0 | probably damaging | 1 | Deleterious | -5.676 | Effect | 64 |
| R755H | Affect | 0 | probably damaging | 1 | Deleterious | -4.822 | Effect | 62 |
| R755C | Affect | 0 | probably damaging | 1 | Deleterious | -7.697 | Effect | 62 |
| V752M | Affect | 0 | probably damaging | 1 | Deleterious | -2.909 | Effect | 50 |
| Y750S | Affect | 0 | probably damaging | 1 | Deleterious | -8.797 | Effect | 81 |
| Y682C | Affect | 0 | probably damaging | 1 | Deleterious | -7.038 | Effect | 44 |
| L606F | Affect | 0 | probably damaging | 1 | Deleterious | -3.44 | Effect | 39 |
| D598N | Affect | 0 | probably damaging | 1 | Deleterious | -4.922 | Effect | 62 |
| G587R | Affect | 0 | probably damaging | 1 | Deleterious | -7.374 | Effect | 56 |
| C539G | Affect | 0 | probably damaging | 1 | Deleterious | -11.611 | Effect | 83 |
| G493R | Affect | 0 | probably damaging | 1 | Deleterious | -7.078 | Effect | 79 |
| L482P | Affect | 0 | probably damaging | 1 | Deleterious | -6.805 | Effect | 89 |
| G452R | Affect | 0 | probably damaging | 1 | Deleterious | -5.917 | Effect | 68 |
| F451L | Affect | 0 | probably damaging | 1 | Deleterious | -5.834 | Effect | 69 |
| G411R | Affect | 0 | probably damaging | 1 | Deleterious | -7.27 | Effect | 61 |
| N408K | Affect | 0 | probably damaging | 1 | Deleterious | -5.692 | Effect | 81 |
| G370R | Affect | 0 | probably damaging | 1 | Deleterious | -7.853 | Effect | 50 |
| P346Q | Affect | 0 | probably damaging | 1 | Deleterious | -5.699 | Effect | 17 |
| P333R | Affect | 0 | probably damaging | 1 | Deleterious | -8.83 | Effect | 42 |
| N316S | Affect | 0 | probably damaging | 1 | Deleterious | -4.906 | Effect | 74 |
| G308D | Affect | 0 | probably damaging | 1 | Deleterious | -6.868 | Effect | 60 |
| E304K | Affect | 0 | probably damaging | 1 | Deleterious | -2.79 | Effect | 10 |
| L297V | Affect | 0 | probably damaging | 1 | Deleterious | -2.743 | Effect | 31 |
| W276R | Affect | 0 | probably damaging | 1 | Deleterious | -13.728 | Effect | 71 |
| C264Y | Affect | 0 | probably damaging | 1 | Deleterious | -10.779 | Effect | 78 |
| P240L | Affect | 0 | probably damaging | 1 | Deleterious | -9.173 | Effect | 26 |
| R184Q | Affect | 0 | probably damaging | 1 | Deleterious | -3.922 | Effect | 60 |
| A123T | Affect | 0 | probably damaging | 1 | Deleterious | -3.911 | Effect | 18 |
| A116G | Affect | 0 | probably damaging | 1 | Deleterious | -3.868 | Effect | 62 |
| C57Y | Affect | 0 | probably damaging | 1 | Deleterious | -10.604 | Effect | 79 |

**Table2:** Disease effect nsSNPs associated variations predicted by SNPs&GO and PHD-SNP softwares:

| dbSNP rs# | SUB | PhD-SNP |  |  | SNPs&GO |  |  |
| --- | --- | --- | --- | --- | --- | --- | --- |
|  |  | prediction | RI | Score | prediction | RI | Score |
| rs1064796541 | E1175K | Disease | 4 | 0.705 | Disease | 6 | 0.822 |
| rs1391080419 | K1150R | Disease | 2 | 0.614 | Disease | 5 | 0.737 |
| rs200590243 | G1149R | Disease | 5 | 0.745 | Disease | 5 | 0.745 |
| rs952497509 | R1145H | Disease | 6 | 0.799 | Disease | 4 | 0.682 |
| rs200798819 | R1145C | Disease | 8 | 0.883 | Disease | 5 | 0.746 |
| rs1363914541 | L1132P | Disease | 7 | 0.838 | Disease | 2 | 0.597 |
| rs782573655 | R990C | Disease | 1 | 0.561 | Disease | 3 | 0.653 |
| rs1375240951 | N945I | Disease | 5 | 0.759 | Disease | 5 | 0.753 |
| rs1273221058 | N825K | Disease | 2 | 0.602 | Disease | 1 | 0.551 |
| rs782502555 | N825Y | Disease | 3 | 0.67 | Disease | 3 | 0.673 |
| rs782776836 | P816S | Disease | 5 | 0.744 | Disease | 4 | 0.724 |
| rs797045674 | Y784C | Disease | 7 | 0.841 | Disease | 2 | 0.622 |
| rs948680832 | R755C | Disease | 3 | 0.636 | Disease | 3 | 0.662 |
| rs1064794855 | Y750S | Disease | 5 | 0.768 | Disease | 3 | 0.661 |
| rs1412295827 | Y682C | Disease | 3 | 0.644 | Disease | 5 | 0.755 |
| rs137852519 | D598N | Disease | 0 | 0.502 | Disease | 4 | 0.676 |
| rs199888009 | G587R | Disease | 6 | 0.825 | Disease | 3 | 0.644 |
| rs886041102 | C539G | Disease | 8 | 0.883 | Disease | 4 | 0.69 |
| rs1312463945 | G493R | Disease | 9 | 0.931 | Disease | 4 | 0.721 |
| rs1064794246 | L482P | Disease | 9 | 0.939 | Disease | 6 | 0.779 |
| rs137852520 | G452R | Disease | 9 | 0.937 | Disease | 6 | 0.784 |
| rs1262433336 | G411R | Disease | 8 | 0.903 | Disease | 3 | 0.661 |
| rs994675918 | N408K | Disease | 9 | 0.942 | Disease | 6 | 0.791 |
| rs137852524 | G370R | Disease | 7 | 0.84 | Disease | 4 | 0.691 |
| rs1064793162 | P333R | Disease | 7 | 0.871 | Disease | 3 | 0.669 |
| rs782052935 | N316S | Disease | 3 | 0.653 | Disease | 3 | 0.654 |
| rs1322838292 | L297V | Disease | 5 | 0.753 | Disease | 3 | 0.661 |
| rs1131691900 | W276R | Disease | 8 | 0.918 | Disease | 5 | 0.731 |
| rs137852518 | C264Y | Disease | 9 | 0.949 | Disease | 6 | 0.785 |
| rs137852526 | P240L | Disease | 6 | 0.788 | Disease | 2 | 0.577 |
| rs137852521 | R184Q | Disease | 6 | 0.791 | Disease | 6 | 0.78 |
| rs782752037 | A123T | Disease | 4 | 0.712 | Disease | 5 | 0.749 |
| rs782704314 | A116G | Disease | 8 | 0.875 | Disease | 4 | 0.689 |
| rs1057517755 | C57Y | Disease | 9 | 0.933 | Disease | 7 | 0.841 |

**\*RI: Reliability Index**

**Table3:**
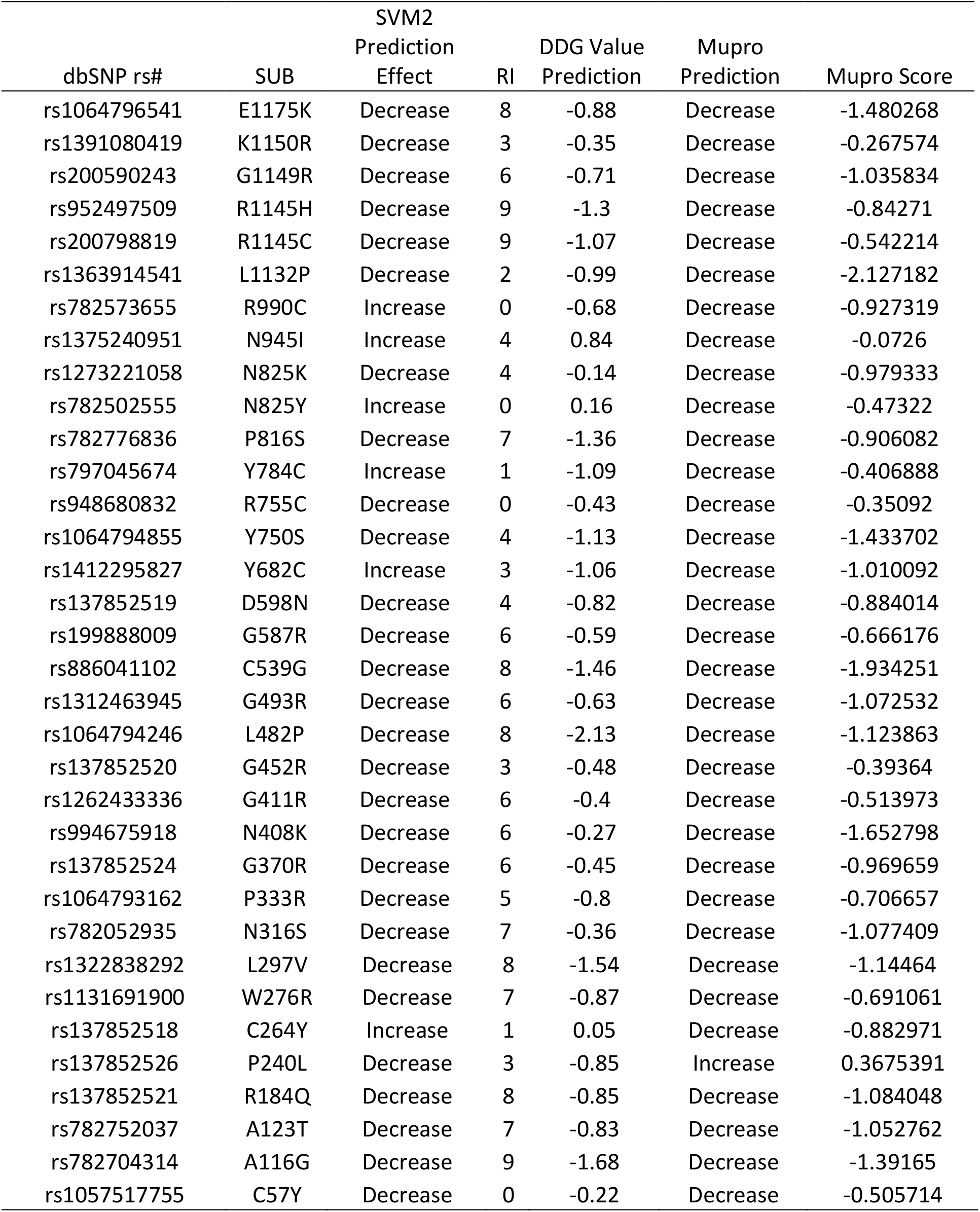
Protein stability analysis predicted by I-Mutant version 3.0 and MUPro:

Interestingly, GeneMANIA could not predict *L1CAM* gene function after the mutations. The genes co-expressed with, share similar protein domain, or participate to achieve similar function were illustrated by GeneMANIA and shown in (figure 2), Tables (4 & 5).

**Figure (2):**
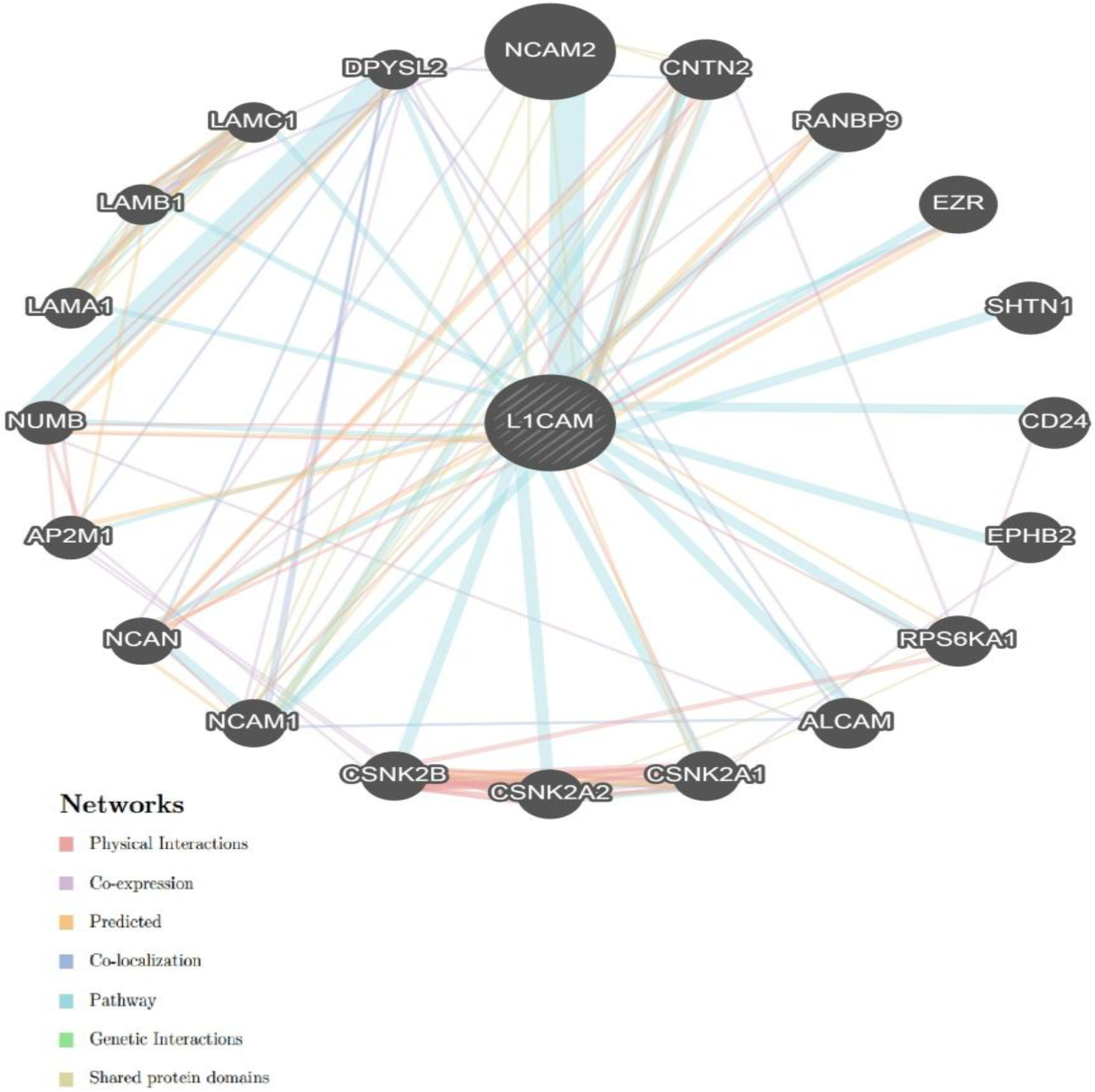
Interaction between *L1CAM* and its related genes.

**Table 4:** The *L1CAM gene* functions and its appearance in network and genome:

| Function | FDR | Genes in network | Genes in genome |
| --- | --- | --- | --- |
| basal lamina | 0.00455213 | 3 | 12 |
| extracellular matrix organization | 0.04748433 | 5 | 292 |
| extracellular structure organization | 0.04748433 | 5 | 293 |
| basement membrane | 0.04748433 | 3 | 33 |
| PcG protein complex | 0.04748433 | 3 | 39 |
| extracellular matrix structural constituent | 0.07853391 | 3 | 53 |
| telencephalon development | 0.10284851 | 3 | 61 |
| extracellular matrix part | 0.11929111 | 3 | 67 |
| cell-cell adhesion | 0.32145746 | 4 | 258 |
| proteinaceous extracellular matrix | 0.41738063 | 3 | 110 |

**\*FDR:** false discovery rate is greater than or equal to the probability that this is a false positive.

**Table (5).**
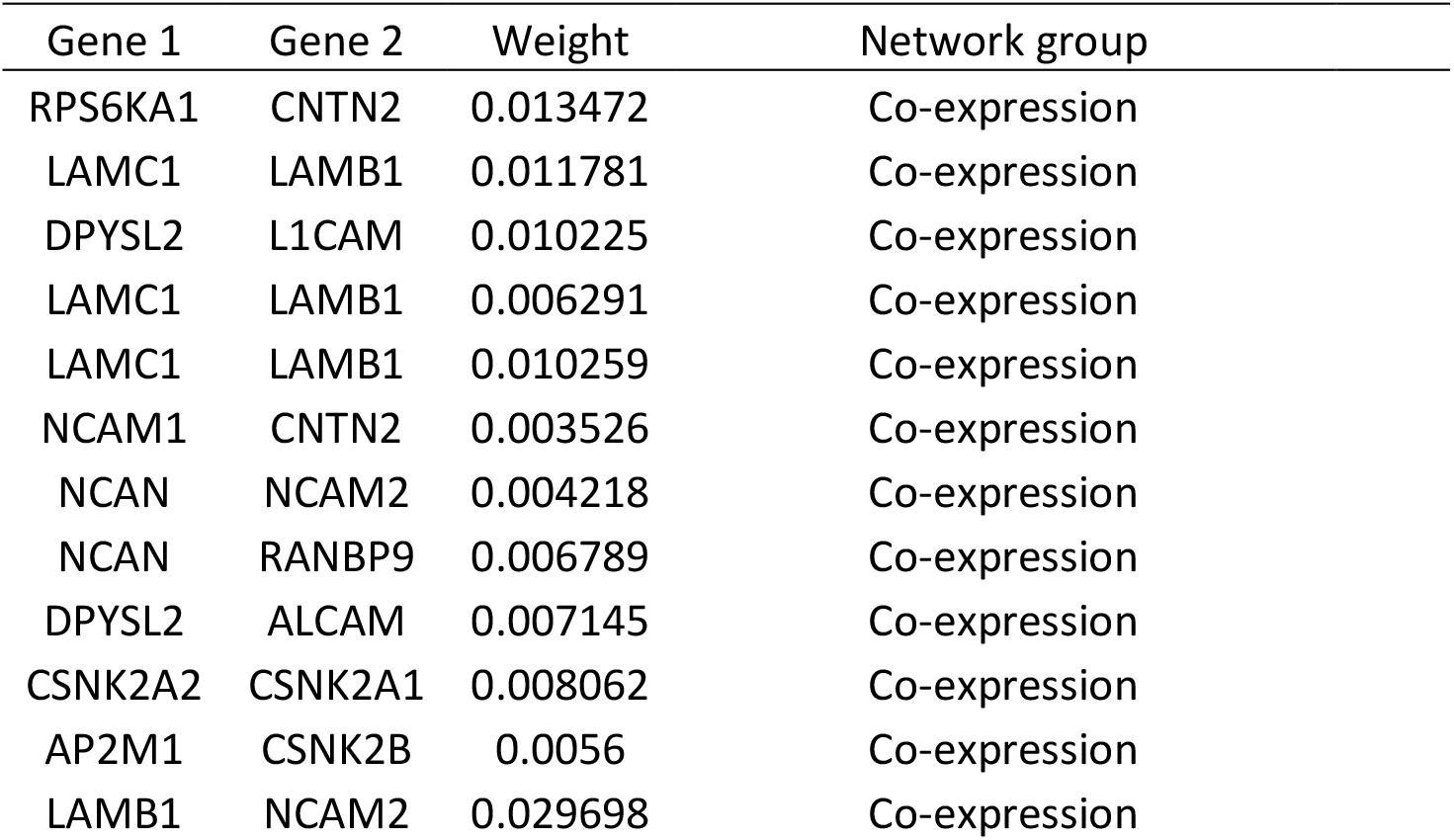
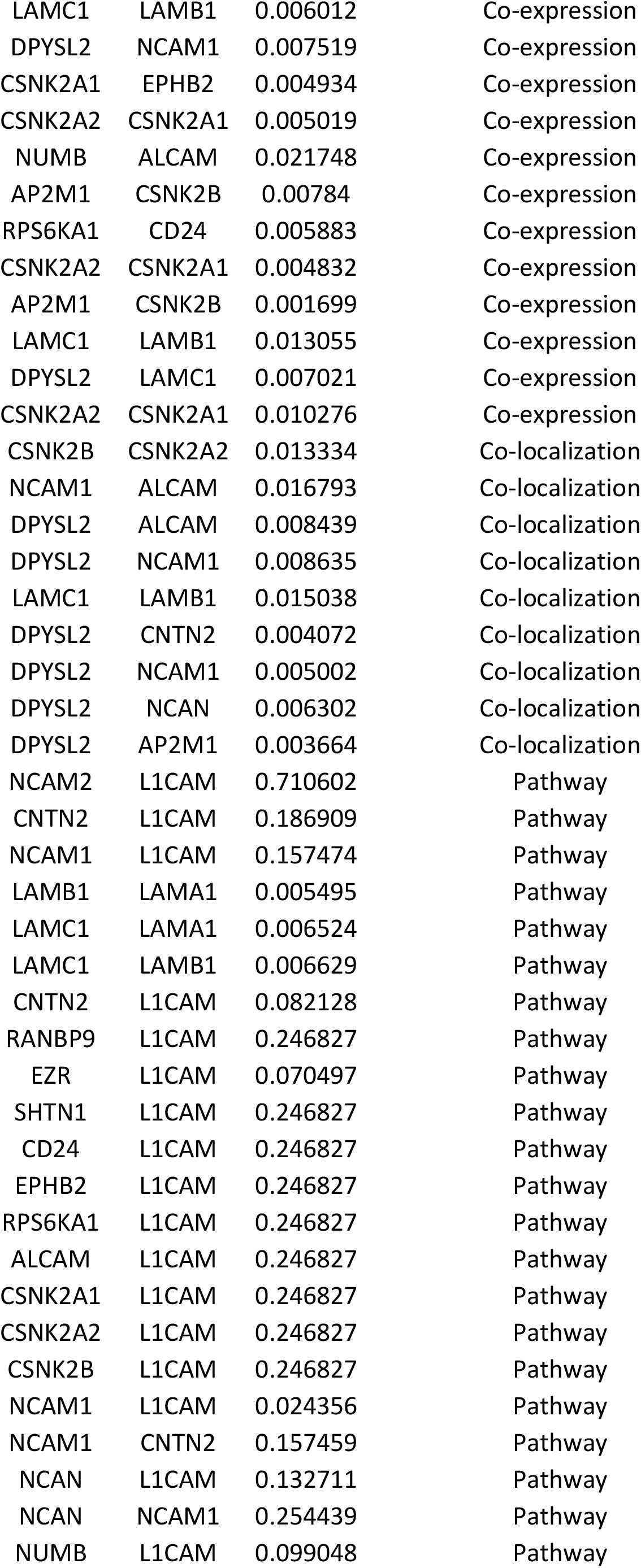
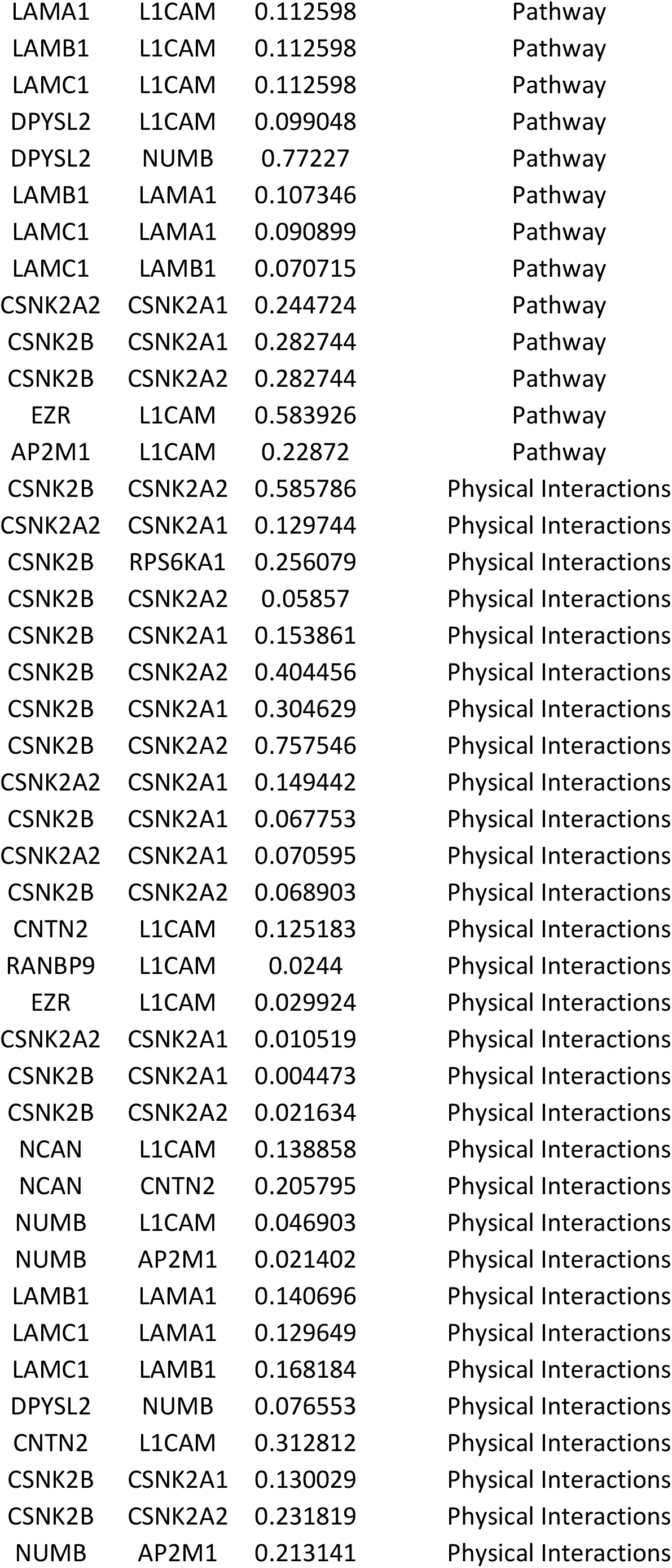
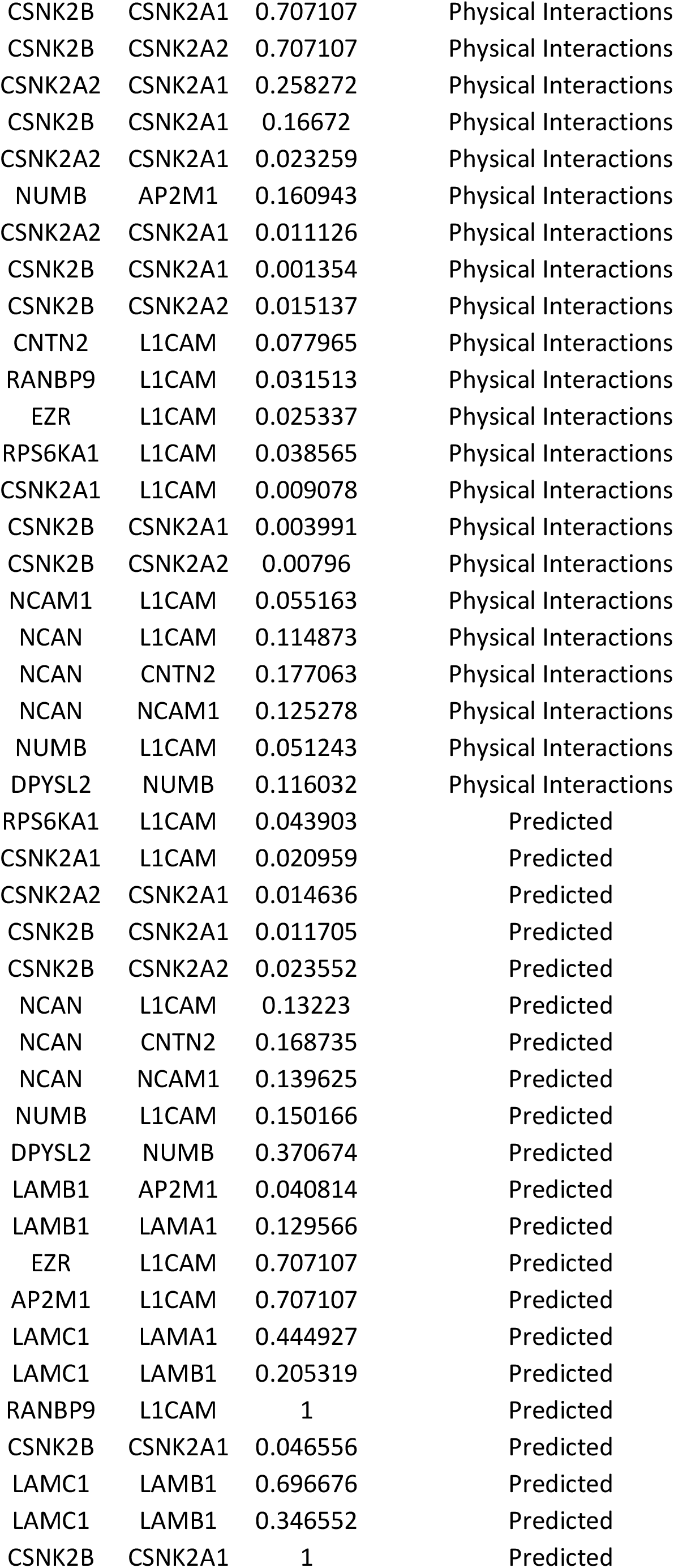
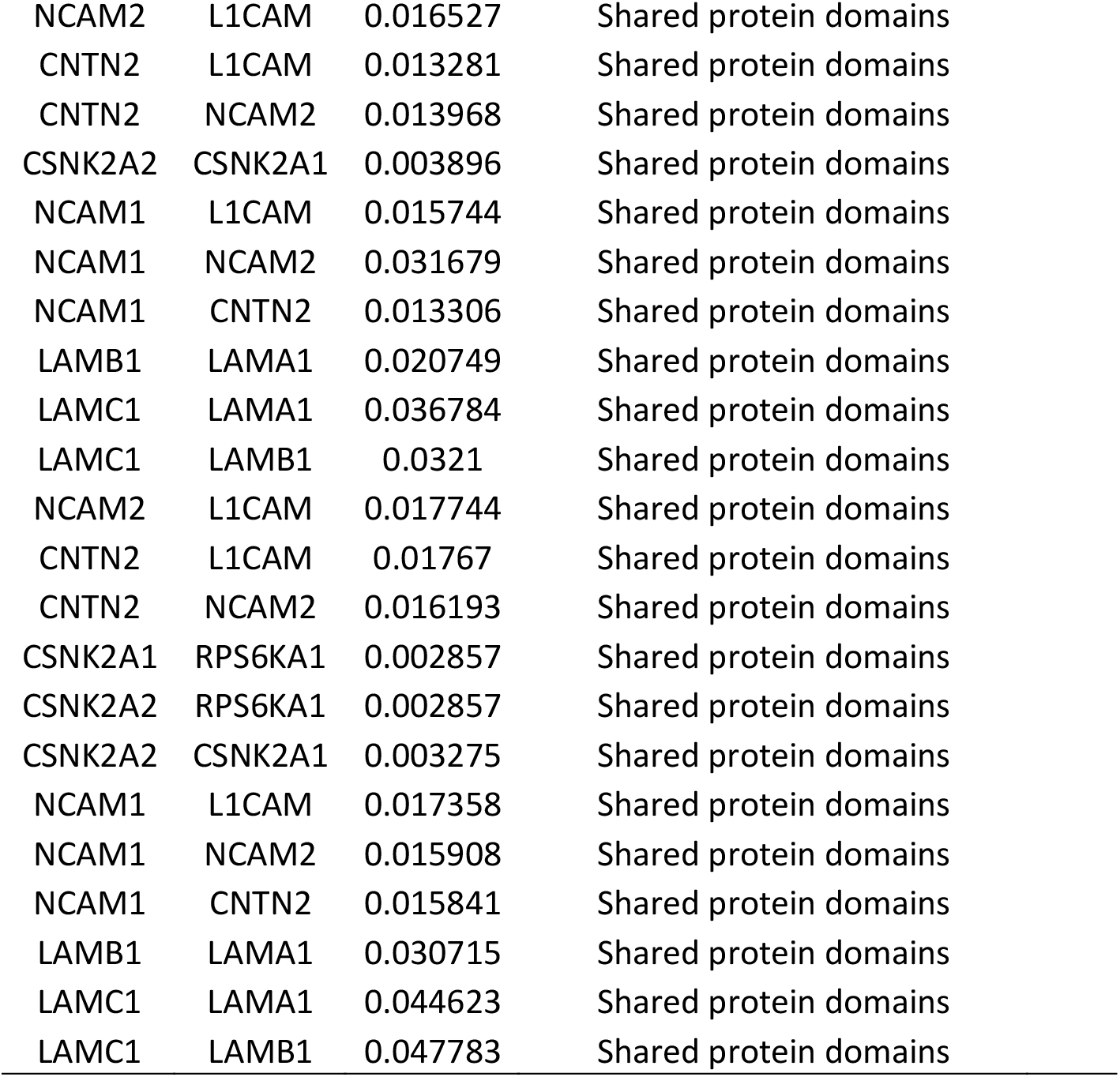
The gene co-expressed, share domain and Interaction with *L1CAM gene* network:

**\*FDR:** false discovery rate is greater than or equal to the probability that this is a false positive. **Table (5) The gene co-expressed, share domain and Interaction with *L1CAM* gene network:**
| Gene 1 | Gene 2 | Weight | Network group |
| --- | --- | --- | --- |
| RPS6KA1 | CNTN2 | 0.013472 | Co-expression |
| LAMC1 | LAMB1 | 0.011781 | Co-expression |
| DPYSL2 | L1CAM | 0.010225 | Co-expression |
| LAMC1 | LAMB1 | 0.006291 | Co-expression |
| LAMC1 | LAMB1 | 0.010259 | Co-expression |
| NCAM1 | CNTN2 | 0.003526 | Co-expression |
| NCAN | NCAM2 | 0.004218 | Co-expression |
| NCAN | RANBP9 | 0.006789 | Co-expression |
| DPYSL2 | ALCAM | 0.007145 | Co-expression |
| CSNK2A2 | CSNK2A1 | 0.008062 | Co-expression |
| AP2M1 | CSNK2B | 0.0056 | Co-expression |
| LAMB1 | NCAM2 | 0.029698 | Co-expression |
| LAMC1 | LAMB1 | 0.006012 | Co-expression |
| DPYSL2 | NCAM1 | 0.007519 | Co-expression |
| CSNK2A1 | EPHB2 | 0.004934 | Co-expression |
| CSNK2A2 | CSNK2A1 | 0.005019 | Co-expression |
| NUMB | ALCAM | 0.021748 | Co-expression |
| AP2M1 | CSNK2B | 0.00784 | Co-expression |
| RPS6KA1 | CD24 | 0.005883 | Co-expression |
| CSNK2A2 | CSNK2A1 | 0.004832 | Co-expression |
| AP2M1 | CSNK2B | 0.001699 | Co-expression |
| LAMC1 | LAMB1 | 0.013055 | Co-expression |
| DPYSL2 | LAMC1 | 0.007021 | Co-expression |
| CSNK2A2 | CSNK2A1 | 0.010276 | Co-expression |
| CSNK2B | CSNK2A2 | 0.013334 | Co-localization |
| NCAM1 | ALCAM | 0.016793 | Co-localization |
| DPYSL2 | ALCAM | 0.008439 | Co-localization |
| DPYSL2 | NCAM1 | 0.008635 | Co-localization |
| LAMC1 | LAMB1 | 0.015038 | Co-localization |
| DPYSL2 | CNTN2 | 0.004072 | Co-localization |
| DPYSL2 | NCAM1 | 0.005002 | Co-localization |
| DPYSL2 | NCAN | 0.006302 | Co-localization |
| DPYSL2 | AP2M1 | 0.003664 | Co-localization |
| NCAM2 | L1CAM | 0.710602 | Pathway |
| CNTN2 | L1CAM | 0.186909 | Pathway |
| NCAM1 | L1CAM | 0.157474 | Pathway |
| LAMB1 | LAMA1 | 0.005495 | Pathway |
| LAMC1 | LAMA1 | 0.006524 | Pathway |
| LAMC1 | LAMB1 | 0.006629 | Pathway |
| CNTN2 | L1CAM | 0.082128 | Pathway |
| RANBP9 | L1CAM | 0.246827 | Pathway |
| EZR | L1CAM | 0.070497 | Pathway |
| SHTN1 | L1CAM | 0.246827 | Pathway |
| CD24 | L1CAM | 0.246827 | Pathway |
| EPHB2 | L1CAM | 0.246827 | Pathway |
| RPS6KA1 | L1CAM | 0.246827 | Pathway |
| ALCAM | L1CAM | 0.246827 | Pathway |
| CSNK2A1 | L1CAM | 0.246827 | Pathway |
| CSNK2A2 | L1CAM | 0.246827 | Pathway |
| CSNK2B | L1CAM | 0.246827 | Pathway |
| NCAM1 | L1CAM | 0.024356 | Pathway |
| NCAM1 | CNTN2 | 0.157459 | Pathway |
| NCAN | L1CAM | 0.132711 | Pathway |
| NCAN | NCAM1 | 0.254439 | Pathway |
| NUMB | L1CAM | 0.099048 | Pathway |
| LAMA1 | L1CAM | 0.112598 | Pathway |
| LAMB1 | L1CAM | 0.112598 | Pathway |
| LAMC1 | L1CAM | 0.112598 | Pathway |
| DPYSL2 | L1CAM | 0.099048 | Pathway |
| DPYSL2 | NUMB | 0.77227 | Pathway |
| LAMB1 | LAMA1 | 0.107346 | Pathway |
| LAMC1 | LAMA1 | 0.090899 | Pathway |
| LAMC1 | LAMB1 | 0.070715 | Pathway |
| CSNK2A2 | CSNK2A1 | 0.244724 | Pathway |
| CSNK2B | CSNK2A1 | 0.282744 | Pathway |
| CSNK2B | CSNK2A2 | 0.282744 | Pathway |
| EZR | L1CAM | 0.583926 | Pathway |
| AP2M1 | L1CAM | 0.22872 | Pathway |
| CSNK2B | CSNK2A2 | 0.585786 | Physical Interactions |
| CSNK2A2 | CSNK2A1 | 0.129744 | Physical Interactions |
| CSNK2B | RPS6KA1 | 0.256079 | Physical Interactions |
| CSNK2B | CSNK2A2 | 0.05857 | Physical Interactions |
| CSNK2B | CSNK2A1 | 0.153861 | Physical Interactions |
| CSNK2B | CSNK2A2 | 0.404456 | Physical Interactions |
| CSNK2B | CSNK2A1 | 0.304629 | Physical Interactions |
| CSNK2B | CSNK2A2 | 0.757546 | Physical Interactions |
| CSNK2A2 | CSNK2A1 | 0.149442 | Physical Interactions |
| CSNK2B | CSNK2A1 | 0.067753 | Physical Interactions |
| CSNK2A2 | CSNK2A1 | 0.070595 | Physical Interactions |
| CSNK2B | CSNK2A2 | 0.068903 | Physical Interactions |
| CNTN2 | L1CAM | 0.125183 | Physical Interactions |
| RANBP9 | L1CAM | 0.0244 | Physical Interactions |
| EZR | L1CAM | 0.029924 | Physical Interactions |
| CSNK2A2 | CSNK2A1 | 0.010519 | Physical Interactions |
| CSNK2B | CSNK2A1 | 0.004473 | Physical Interactions |
| CSNK2B | CSNK2A2 | 0.021634 | Physical Interactions |
| NCAN | L1CAM | 0.138858 | Physical Interactions |
| NCAN | CNTN2 | 0.205795 | Physical Interactions |
| NUMB | L1CAM | 0.046903 | Physical Interactions |
| NUMB | AP2M1 | 0.021402 | Physical Interactions |
| LAMB1 | LAMA1 | 0.140696 | Physical Interactions |
| LAMC1 | LAMA1 | 0.129649 | Physical Interactions |
| LAMC1 | LAMB1 | 0.168184 | Physical Interactions |
| DPYSL2 | NUMB | 0.076553 | Physical Interactions |
| CNTN2 | L1CAM | 0.312812 | Physical Interactions |
| CSNK2B | CSNK2A1 | 0.130029 | Physical Interactions |
| CSNK2B | CSNK2A2 | 0.231819 | Physical Interactions |
| NUMB | AP2M1 | 0.213141 | Physical Interactions |
| CSNK2B | CSNK2A1 | 0.707107 | Physical Interactions |
| CSNK2B | CSNK2A2 | 0.707107 | Physical Interactions |
| CSNK2A2 | CSNK2A1 | 0.258272 | Physical Interactions |
| CSNK2B | CSNK2A1 | 0.16672 | Physical Interactions |
| CSNK2A2 | CSNK2A1 | 0.023259 | Physical Interactions |
| NUMB | AP2M1 | 0.160943 | Physical Interactions |
| CSNK2A2 | CSNK2A1 | 0.011126 | Physical Interactions |
| CSNK2B | CSNK2A1 | 0.001354 | Physical Interactions |
| CSNK2B | CSNK2A2 | 0.015137 | Physical Interactions |
| CNTN2 | L1CAM | 0.077965 | Physical Interactions |
| RANBP9 | L1CAM | 0.031513 | Physical Interactions |
| EZR | L1CAM | 0.025337 | Physical Interactions |
| RPS6KA1 | L1CAM | 0.038565 | Physical Interactions |
| CSNK2A1 | L1CAM | 0.009078 | Physical Interactions |
| CSNK2B | CSNK2A1 | 0.003991 | Physical Interactions |
| CSNK2B | CSNK2A2 | 0.00796 | Physical Interactions |
| NCAM1 | L1CAM | 0.055163 | Physical Interactions |
| NCAN | L1CAM | 0.114873 | Physical Interactions |
| NCAN | CNTN2 | 0.177063 | Physical Interactions |
| NCAN | NCAM1 | 0.125278 | Physical Interactions |
| NUMB | L1CAM | 0.051243 | Physical Interactions |
| DPYSL2 | NUMB | 0.116032 | Physical Interactions |
| RPS6KA1 | L1CAM | 0.043903 | Predicted |
| CSNK2A1 | L1CAM | 0.020959 | Predicted |
| CSNK2A2 | CSNK2A1 | 0.014636 | Predicted |
| CSNK2B | CSNK2A1 | 0.011705 | Predicted |
| CSNK2B | CSNK2A2 | 0.023552 | Predicted |
| NCAN | L1CAM | 0.13223 | Predicted |
| NCAN | CNTN2 | 0.168735 | Predicted |
| NCAN | NCAM1 | 0.139625 | Predicted |
| NUMB | L1CAM | 0.150166 | Predicted |
| DPYSL2 | NUMB | 0.370674 | Predicted |
| LAMB1 | AP2M1 | 0.040814 | Predicted |
| LAMB1 | LAMA1 | 0.129566 | Predicted |
| EZR | L1CAM | 0.707107 | Predicted |
| AP2M1 | L1CAM | 0.707107 | Predicted |
| LAMC1 | LAMA1 | 0.444927 | Predicted |
| LAMC1 | LAMB1 | 0.205319 | Predicted |
| RANBP9 | L1CAM | 1 | Predicted |
| CSNK2B | CSNK2A1 | 0.046556 | Predicted |
| LAMC1 | LAMB1 | 0.696676 | Predicted |
| LAMC1 | LAMB1 | 0.346552 | Predicted |
| CSNK2B | CSNK2A1 | 1 | Predicted |
| NCAM2 | L1CAM | 0.016527 | Shared protein domains |
| CNTN2 | L1CAM | 0.013281 | Shared protein domains |
| CNTN2 | NCAM2 | 0.013968 | Shared protein domains |
| CSNK2A2 | CSNK2A1 | 0.003896 | Shared protein domains |
| NCAM1 | L1CAM | 0.015744 | Shared protein domains |
| NCAM1 | NCAM2 | 0.031679 | Shared protein domains |
| NCAM1 | CNTN2 | 0.013306 | Shared protein domains |
| LAMB1 | LAMA1 | 0.020749 | Shared protein domains |
| LAMC1 | LAMA1 | 0.036784 | Shared protein domains |
| LAMC1 | LAMB1 | 0.0321 | Shared protein domains |
| NCAM2 | L1CAM | 0.017744 | Shared protein domains |
| CNTN2 | L1CAM | 0.01767 | Shared protein domains |
| CNTN2 | NCAM2 | 0.016193 | Shared protein domains |
| CSNK2A1 | RPS6KA1 | 0.002857 | Shared protein domains |
| CSNK2A2 | RPS6KA1 | 0.002857 | Shared protein domains |
| CSNK2A2 | CSNK2A1 | 0.003275 | Shared protein domains |
| NCAM1 | L1CAM | 0.017358 | Shared protein domains |
| NCAM1 | NCAM2 | 0.015908 | Shared protein domains |
| NCAM1 | CNTN2 | 0.015841 | Shared protein domains |
| LAMB1 | LAMA1 | 0.030715 | Shared protein domains |
| LAMC1 | LAMA1 | 0.044623 | Shared protein domains |
| LAMC1 | LAMB1 | 0.047783 | Shared protein domains |

**Table (6):**
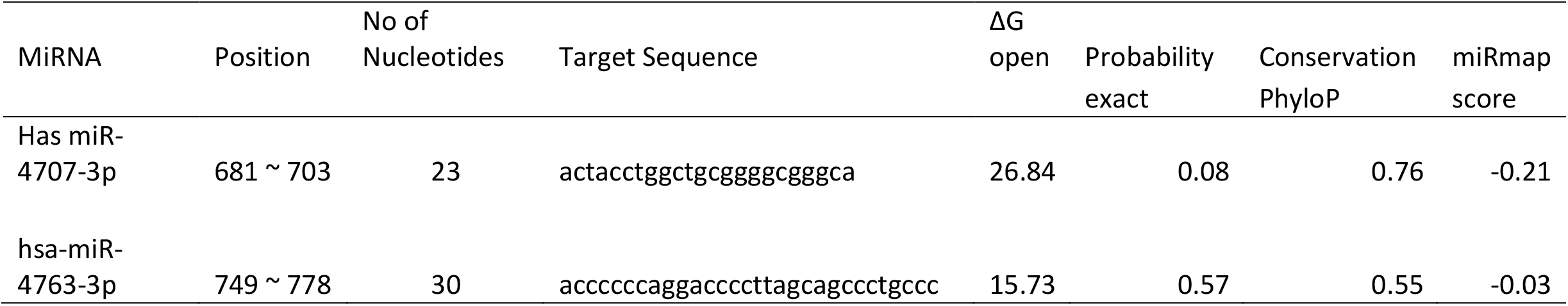
Common miRNAs identified in 3 UTR region by miRmap software:

**Table (7):**
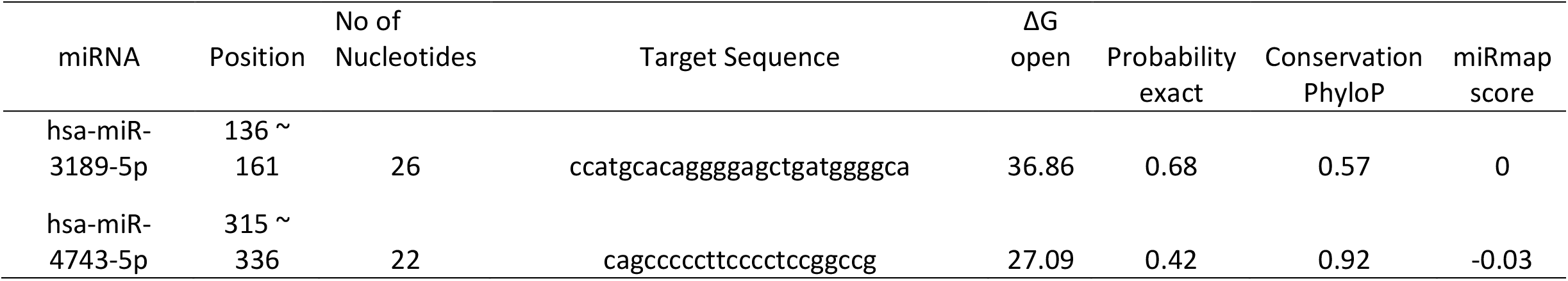
Common miRNAs identified in 5 UTR region by miRmap software:

We also used ConSurf server; the nsSNPs that are located at highly conserved amino acid positions tend to be more deleterious than nsSNPs that are located at non-conserved sites. (supplemental figure 4 for ConSurf result, which is available at https://www.biorxiv.org/)

There were some studies that have been reported which show pathogenic nsSNPs that cause L1 syndrome (11, 70-78), which is corresponding with our results. It has been observed that *L1CAM* polymorphisms are associated in several types of human cancers (27-50, 79). Furthermore, this study confirm that (E1175K, Y784C, Y750S, D598N, C539G, G452R, G370R, P333R, W276R, C264Y, P240L and R184Q) SNPs are pathogenic, these results show similarities with the results found earlier in dbSNPs/NCBI database. Also all these SNPs were retrieved as untested (C57Y, A116G, A123T, L297V, N316S, N408K, G411R, G493R, Y682C, R755C, P816S, N825Y, N825K, N945I, R990C, L1132P, R1145C, R1145H, G1149R, and K1150R) were found to be all pathogenic.

We also used Mutation3D server; (Figure 3) All SNPs in red (R184Q, P240L, C264Y, W276R, L297V and N316S) are clustered mutation. Significantly, such mutation clusters are abundant in human cancers (82) we think it is associated with breast cancer. Afflictions which can be considered cases of highly accelerated evolution within somatic tissues. Recent studies have revealed several molecular mechanisms of clustered mutagenesis. (83) While SNPs in blue (C57Y, A116G, A123T, P333R, G370R, N408K and G411R) and gray are covered and uncovered mutation respectively.

**Figure (3):**
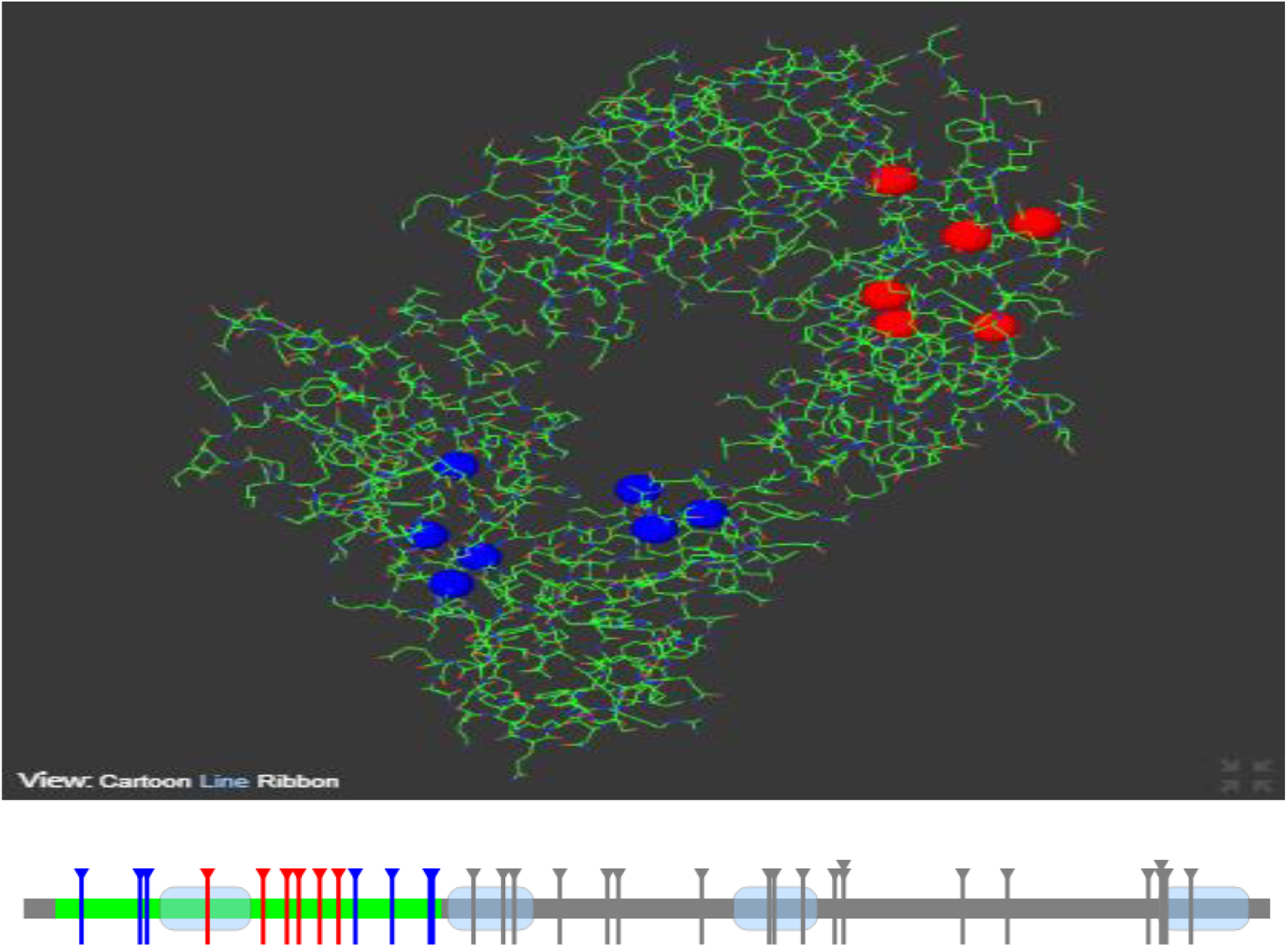
Structural models for wild type *L1CAM*, illustrated by Mutation3D.

Different cellular functions are controlled by transcriptional regulation done by non-coding RNA molecules. Recently discovered class of non-coding RNA molecules is MicroRNA (miRNA) that are small non-coding RNA having function of activation and/or suppression of protein translation inside the cells at post-transcriptional level (84). In present study, we have predicted some targets miRNAs for *L1CAM* gene. we found that it has sites for (Has miR-4707-3p) and (hsa-miR-4763-3p) microRNA in the 3 UTR regions. And sites for (hsa-miR-3189-5p) and (hsa-miR-4743-5p) microRNA in the 5 regions. Which were noted to be conserved among different species indicating its significant role in the function of the final protein. This insight provides clue to wet-lab researches to understand the expression pattern of *L1CAM* gene and binding phenomenon of mRNA and miRNA upon mutation.

This study revealed *26 novel* Pathological mutations that have a potential functional impact and may thus be used as diagnostic markers for L1 syndrome and can make an ideal target for tumor therapy (32, 80) properties of *L1CAM*, in addition to its cell surface localization, make it a potentially useful diagnostic marker for cancer progression and a candidate for anti-cancer therapy (39, 81). Finally some appreciations of wet lab techniques are suggested to support these findings.

## 5. Conclusion

A large number of different pathological *L1CAM* mutations have been identified. Therefore the confirmation of these nsSNPs in L1 syndrome was crucial by using Comprehensive bioinformatics analysis. These findings describe *26* novel L1 mutations which improve our understanding on genotype-phenotype correlation. And can be used as diagnostic markers for L1 syndrome and besides in cancer diagnosis.

## Supporting information

supplemental figure 4 for ConSurf result

## Conflict of interest

The authors declare no conflict of interest.

## Acknowledgment

The authors wish to acknowledgment the enthusiastic cooperation of Africa City of Technology -Sudan.

## References

1. Isik E, Onay H, Atik T, Akgun B, Cogulu O, Ozkinay F. Clinical and genetic features of L1 syndrome patients: Definition of two novel mutations. Clinical neurology and neurosurgery. 2018;172:20–3.

2. Zhang L. CRASH syndrome: does it teach us about neurotrophic functions of cell adhesion molecules? The Neuroscientist: a review journal bringing neurobiology, neurology and psychiatry. 2010;16(4):470–4.

3. Chidsey BA, Baldwin EE, Toydemir R, Ahles L, Hanson H, Stevenson DA. L1CAM whole gene deletion in a child with L1 syndrome. American journal of medical genetics Part A. 2014;164a(6):1555–8.

4. Ferese R, Zampatti S, Griguoli AM, Fornai F, Giardina E, Barrano G, et al. A New Splicing Mutation in the L1CAM Gene Responsible for X-Linked Hydrocephalus (HSAS). Journal of molecular neuroscience: MN. 2016;59(3):376–81.

5. Fernandez RM, Nunez-Torres R, Garcia-Diaz L, de Agustin JC, Antinolo G, Borrego S. Association of X-linked hydrocephalus and Hirschsprung disease: report of a new patient with a mutation in the L1CAM gene. American journal of medical genetics Part A. 2012;158a(4):816–20.

6. Weller S, Gartner J. Genetic and clinical aspects of X-linked hydrocephalus (L1 disease): Mutations in the L1CAM gene. Human mutation. 2001;18(1):1–12.

7. Shaw M, Yap TY, Henden L, Bahlo M, Gardner A, Kalscheuer VM, et al. Identical by descent L1CAM mutation in two apparently unrelated families with intellectual disability without L1 syndrome. European journal of medical genetics. 2015;58(6-7):364–8.

8. Sztriha L, Frossard P, Hofstra RM, Verlind E, Nork M. Novel missense mutation in the L1 gene in a child with corpus callosum agenesis, retardation, adducted thumbs, spastic paraparesis, and hydrocephalus. Journal of child neurology. 2000;15(4):239–43.

9. Panayi M, Gokhale D, Mansour S, Elles R. Prenatal diagnosis in a family with X-linked hydrocephalus. Prenatal diagnosis. 2005;25(10):930–3.

10. Ochando I, Vidal V, Gascon J, Acien M, Urbano A, Rueda J. Prenatal diagnosis of X-linked hydrocephalus in a family with a novel mutation in L1CAM gene. Journal of obstetrics and gynaecology: the journal of the Institute of Obstetrics and Gynaecology. 2016;36(3):403–5.

11. Finckh U, Schroder J, Ressler B, Veske A, Gal A. Spectrum and detection rate of L1CAM mutations in isolated and familial cases with clinically suspected L1-disease. American journal of medical genetics. 2000;92(1):40–6.

12. Fransen E, Van Camp G, D’Hooge R, Vits L, Willems PJ. Genotype-phenotype correlation in L1 associated diseases. Journal of medical genetics. 1998;35(5):399–404.

13. Christaller WA, Vos Y, Gebre-Medhin S, Hofstra RM, Schafer MK. L1 syndrome diagnosis complemented with functional analysis of L1CAM variants located to the two N-terminal Ig-like domains. Clinical genetics. 2017;91(1):115–20.

14. Yamasaki M, Thompson P, Lemmon V. CRASH syndrome: mutations in L1CAM correlate with severity of the disease. Neuropediatrics. 1997;28(3):175–8.

15. Bertolin C, Boaretto F, Barbon G, Salviati L, Lapi E, Divizia MT, et al. Novel mutations in the L1CAM gene support the complexity of L1 syndrome. Journal of the neurological sciences. 2010;294(1-2):124–6.

16. Vits L, Van Camp G, Coucke P, Fransen E, De Boulle K, Reyniers E, et al. MASA syndrome is due to mutations in the neural cell adhesion gene L1CAM. Nature genetics. 1994;7(3):408–13.

17. Vos YJ, Hofstra RM. An updated and upgraded L1CAM mutation database. Human mutation. 2010;31(1):E1102–9.

18. Simonati A, Boaretto F, Vettori A, Dabrilli P, Criscuolo L, Rizzuto N, et al. A novel missense mutation in the L1CAM gene in a boy with L1 disease. Neurological sciences: official journal of the Italian Neurological Society and of the Italian Society of Clinical Neurophysiology. 2006;27(2):114–7.

19. Marin R, Ley-Martos M, Gutierrez G, Rodriguez-Sanchez F, Arroyo D, Mora-Lopez F. Three cases with L1 syndrome and two novel mutations in the L1CAM gene. European journal of pediatrics. 2015;174(11):1541–4.

20. Dietrich A, Korn B, Poustka A. Completion of the physical map of Xq28: the location of the gene for L1CAM on the human X chromosome. Mammalian genome: official journal of the International Mammalian Genome Society. 1992;3(3):168–72.

21. Fransen E, Lemmon V, Van Camp G, Vits L, Coucke P, Willems PJ. CRASH syndrome: clinical spectrum of corpus callosum hypoplasia, retardation, adducted thumbs, spastic paraparesis and hydrocephalus due to mutations in one single gene, L1. European journal of human genetics: EJHG. 1995;3(5):273–84.

22. Jouet M, Feldman E, Yates J, Donnai D, Paterson J, Siggers D, et al. Refining the genetic location of the gene for X linked hydrocephalus within Xq28. Journal of medical genetics. 1993;30(3):214–7.

23. Panicker AK, Buhusi M, Thelen K, Maness PF. Cellular signalling mechanisms of neural cell adhesion molecules. Frontiers in bioscience: a journal and virtual library. 2003;8:d900–11.

24. Sytnyk V, Leshchyns’ka I, Schachner M. Neural Cell Adhesion Molecules of the Immunoglobulin Superfamily Regulate Synapse Formation, Maintenance, and Function. Trends in neurosciences. 2017;40(5):295–308.

25. Itoh K, Fushiki S. The role of L1cam in murine corticogenesis, and the pathogenesis of hydrocephalus. Pathology international. 2015;65(2):58–66.

26. Patzke C, Acuna C, Giam LR, Wernig M, Sudhof TC. Conditional deletion of L1CAM in human neurons impairs both axonal and dendritic arborization and action potential generation. The Journal of experimental medicine. 2016;213(4):499–515.

27. Kiefel H, Bondong S, Hazin J, Ridinger J, Schirmer U, Riedle S, et al. L1CAM: a major driver for tumor cell invasion and motility. Cell adhesion & migration. 2012;6(4):374–84.

28. Lund K, Dembinski JL, Solberg N, Urbanucci A, Mills IG, Krauss S. Slug-dependent upregulation of L1CAM is responsible for the increased invasion potential of pancreatic cancer cells following long-term 5-FU treatment. PloS one. 2015;10(4):e0123684.

29. Haase G, Gavert N, Brabletz T, Ben-Ze’ev A. A point mutation in the extracellular domain of L1 blocks its capacity to confer metastasis in colon cancer cells via CD10. Oncogene. 2017;36(11):1597–606.

30. Samatov TR, Wicklein D, Tonevitsky AG. L1CAM: Cell adhesion and more. Progress in histochemistry and cytochemistry. 2016;51(2):25–32.

31. Schafer MK, Altevogt P. L1CAM malfunction in the nervous system and human carcinomas. Cellular and molecular life sciences: CMLS. 2010;67(14):2425–37.

32. Doberstein K, Milde-Langosch K, Bretz NP, Schirmer U, Harari A, Witzel I, et al. L1CAM is expressed in triple-negative breast cancers and is inversely correlated with androgen receptor. BMC cancer. 2014;14:958.

33. Pechriggl EJ, Concin N, Blumer MJ, Bitsche M, Zwierzina M, Dudas J, et al. L1CAM in the Early Enteric and Urogenital System. The journal of histochemistry and cytochemistry: official journal of the Histochemistry Society. 2017;65(1):21–32.

34. Altevogt P, Doberstein K, Fogel M. L1CAM in human cancer. International journal of cancer. 2016;138(7):1565–76.

35. Guo JC, Xie YM, Ran LQ, Cao HH, Sun C, Wu JY, et al. L1CAM drives oncogenicity in esophageal squamous cell carcinoma by stimulation of ezrin transcription. Journal of molecular medicine (Berlin, Germany). 2017;95(12):1355–68.

36. Klat J, Mladenka A, Dvorackova J, Bajsova S, Simetka O. L1CAM as a Negative Prognostic Factor in Endometrioid Endometrial Adenocarcinoma FIGO Stage IA-IB. Anticancer research. 2019;39(1):421–4.

37. Cavallaro U. L1 in tumor vasculature. Oncotarget. 2016;7(11):11744–5.

38. Kim H, Hwang H, Lee H, Hong HJ. L1 Cell Adhesion Molecule Promotes Migration and Invasion via JNK Activation in Extrahepatic Cholangiocarcinoma Cells with Activating KRAS Mutation. Molecules and cells. 2017;40(5):363–70.

39. Yu X, Yang F, Fu DL, Jin C. L1 cell adhesion molecule as a therapeutic target in cancer. Expert review of anticancer therapy. 2016;16(3):359–71.

40. Ernst AK, Putscher A, Samatov TR, Suling A, Galatenko VV, Shkurnikov MY, et al. Knockdown of L1CAM significantly reduces metastasis in a xenograft model of human melanoma: L1CAM is a potential target for anti-melanoma therapy. PloS one. 2018;13(2):e0192525.

41. Schirmer U, Fiegl H, Pfeifer M, Zeimet AG, Muller-Holzner E, Bode PK, et al. Epigenetic regulation of L1CAM in endometrial carcinoma: comparison to cancer-testis (CT-X) antigens. BMC cancer. 2013;13:156.

42. Sluiter N, de Cuba E, Kwakman R, Kazemier G, Meijer G, Te Velde EA. Adhesion molecules in peritoneal dissemination: function, prognostic relevance and therapeutic options. Clinical & experimental metastasis. 2016;33(5):401–16.

43. Chong Y, Zhang J, Guo X, Li G, Zhang S, Li C, et al. MicroRNA-503 acts as a tumor suppressor in osteosarcoma by targeting L1CAM. PloS one. 2014;9(12):e114585.

44. Sung SY, Wu IH, Chuang PH, Petros JA, Wu HC, Zeng HJ, et al. Targeting L1 cell adhesion molecule expression using liposome-encapsulated siRNA suppresses prostate cancer bone metastasis and growth. Oncotarget. 2014;5(20):9911–29.

45. Manderson EN, Birch AH, Shen Z, Mes-Masson AM, Provencher D, Tonin PN. Molecular genetic analysis of a cell adhesion molecule with homology to L1CAM, contactin 6, and contactin 4 candidate chromosome 3p26pter tumor suppressor genes in ovarian cancer. International journal of gynecological cancer: official journal of the International Gynecological Cancer Society. 2009;19(4):513–25.

46. Notaro S, Reimer D, Duggan-Peer M, Fiegl H, Wiedermair A, Rossler J, et al. Evaluating L1CAM expression in human endometrial cancer using qRT-PCR. Oncotarget. 2016;7(26):40221–32.

47. Kiefel H, Pfeifer M, Bondong S, Hazin J, Altevogt P. Linking L1CAM-mediated signaling to NF-kappaB activation. Trends in molecular medicine. 2011;17(4):178–87.

48. Kiefel H, Bondong S, Pfeifer M, Schirmer U, Erbe-Hoffmann N, Schafer H, et al. EMT-associated up-regulation of L1CAM provides insights into L1CAM-mediated integrin signalling and NF-kappaB activation. Carcinogenesis. 2012;33(10):1919–29.

49. Tischler V, Pfeifer M, Hausladen S, Schirmer U, Bonde AK, Kristiansen G, et al. L1CAM protein expression is associated with poor prognosis in non-small cell lung cancer. Molecular cancer. 2011;10:127.

50. Jo DH, Lee K, Kim JH, Jun HO, Kim Y, Cho YL, et al. L1 increases adhesion-mediated proliferation and chemoresistance of retinoblastoma. Oncotarget. 2017;8(9):15441–52.

51. Adle-Biassette H, Saugier-Veber P, Fallet-Bianco C, Delezoide AL, Razavi F, Drouot N, et al. Neuropathological review of 138 cases genetically tested for X-linked hydrocephalus: evidence for closely related clinical entities of unknown molecular bases. Acta neuropathologica. 2013;126(3):427–42.

52. Stumpel C, Vos YJ. L1 Syndrome. In: Adam MP, Ardinger HH, Pagon RA, Wallace SE, Bean LJH, Stephens K, et al., editors. GeneReviews((R)). Seattle (WA): University of Washington, Seattle University of Washington, Seattle. GeneReviews is a registered trademark of the University of Washington, Seattle. All rights reserved.; 1993.

53. Tenenbaum JD. Translational Bioinformatics: Past, Present, and Future. Genomics, proteomics & bioinformatics. 2016;14(1):31–41.

54. Database resources of the National Center for Biotechnology Information. Nucleic acids research. 2018;46(D1):D8–d13.

55. UniProt: the universal protein knowledgebase. Nucleic acids research. 2017;45(D1):D158–d69.

56. Schneider G, Hu J, Sim N-L, Kumar P, Henikoff S, Ng PC. SIFT web server: predicting effects of amino acid substitutions on proteins. Nucleic acids research. 2012;40(W1):W452–W7.

57. Ng PC, Henikoff S. SIFT: Predicting amino acid changes that affect protein function. Nucleic acids research. 2003;31(13):3812–4.

58. Adzhubei IA, Schmidt S, Peshkin L, Ramensky VE, Gerasimova A, Bork P, et al. A method and server for predicting damaging missense mutations. Nature methods. 2010;7(4):248–9.

59. Ramensky V, Bork P, Sunyaev S. Human non-synonymous SNPs: server and survey. Nucleic acids research. 2002;30(17):3894–900.

60. Choi Y, Sims GE, Murphy S, Miller JR, Chan AP. Predicting the functional effect of amino acid substitutions and indels. PloS one. 2012;7(10):e46688.

61. Choi Y, Chan AP. PROVEAN web server: a tool to predict the functional effect of amino acid substitutions and indels. Bioinformatics. 2015;31(16):2745–7.

62. Capriotti E, Calabrese R, Fariselli P, Martelli PL, Altman RB, Casadio R. WS-SNPs&GO: a web server for predicting the deleterious effect of human protein variants using functional annotation. BMC genomics. 2013;14 Suppl 3:S6.

63. Capriotti E, Martelli PL, Fariselli P, Casadio R. Blind prediction of deleterious amino acid variations with SNPs&GO. Human mutation. 2017;38(9):1064–71.

64. Capriotti E, Fariselli P, Casadio R. I-Mutant2.0: predicting stability changes upon mutation from the protein sequence or structure. Nucleic acids research. 2005;33(Web Server issue):W306–10.

65. Montojo J, Zuberi K, Rodriguez H, Kazi F, Wright G, Donaldson SL, et al. GeneMANIA Cytoscape plugin: fast gene function predictions on the desktop. Bioinformatics. 2010;26(22):2927–8.

66. Warde-Farley D, Donaldson SL, Comes O, Zuberi K, Badrawi R, Chao P, et al. The GeneMANIA prediction server: biological network integration for gene prioritization and predicting gene function. Nucleic acids research. 2010;38(Web Server issue):W214–20.

67. Franz M, Rodriguez H, Lopes C, Zuberi K, Montojo J, Bader GD, et al. GeneMANIA update 2018. Nucleic acids research. 2018;46(W1):W60–w4.

68. Meyer MJ, Lapcevic R, Romero AE, Yoon M, Das J, Beltran JF, et al. mutation3D: Cancer Gene Prediction Through Atomic Clustering of Coding Variants in the Structural Proteome. Human mutation. 2016;37(5):447–56.

69. Shaw G. Polymorphism and single nucleotide polymorphisms (SNPs). BJU international. 2013;112(5):664–5.

70. Marx M, Diestel S, Bozon M, Keglowich L, Drouot N, Bouche E, et al. Pathomechanistic characterization of two exonic L1CAM variants located in trans in an obligate carrier of X-linked hydrocephalus. Neurogenetics. 2012;13(1):49–59.

71. Kudumala S, Freund J, Hortsch M, Godenschwege TA. Differential effects of human L1CAM mutations on complementing guidance and synaptic defects in Drosophila melanogaster. PloS one. 2013;8(10):e76974.

72. Schafer MK, Nam YC, Moumen A, Keglowich L, Bouche E, Kuffner M, et al. L1 syndrome mutations impair neuronal L1 function at different levels by divergent mechanisms. Neurobiology of disease. 2010;40(1):222–37.

73. Randles LG, Lappalainen I, Fowler SB, Moore B, Hamill SJ, Clarke J. Using model proteins to quantify the effects of pathogenic mutations in Ig-like proteins. The Journal of biological chemistry. 2006;281(34):24216–26.

74. Basel-Vanagaite L, Straussberg R, Friez MJ, Inbar D, Korenreich L, Shohat M, et al. Expanding the phenotypic spectrum of L1CAM-associated disease. Clinical genetics. 2006;69(5):414–9.

75. Runker AE, Bartsch U, Nave KA, Schachner M. The C264Y missense mutation in the extracellular domain of L1 impairs protein trafficking in vitro and in vivo. The Journal of neuroscience: the official journal of the Society for Neuroscience. 2003;23(1):277–86.

76. Tagliavacca L, Colombo F, Racchetti G, Meldolesi J. L1CAM and its cell-surface mutants: new mechanisms and effects relevant to the physiology and pathology of neural cells. Journal of neurochemistry. 2013;124(3):397–409.

77. Cheng L, Lemmon V. Pathological missense mutations of neural cell adhesion molecule L1 affect neurite outgrowth and branching on an L1 substrate. Molecular and cellular neurosciences. 2004;27(4):522–30.

78. De Angelis E, Watkins A, Schafer M, Brummendorf T, Kenwrick S. Disease-associated mutations in L1 CAM interfere with ligand interactions and cell-surface expression. Human molecular genetics. 2002;11(1):1–12.

79. Zhang J, Yang F, Ding Y, Zhen L, Han X, Jiao F, et al. Overexpression of L1 cell adhesion molecule correlates with aggressive tumor progression of patients with breast cancer and promotes motility of breast cancer cells. International journal of clinical and experimental pathology. 2015;8(8):9240–7.

80. Gavert N, Ben-Shmuel A, Raveh S, Ben-Ze’ev A. L1-CAM in cancerous tissues. Expert opinion on biological therapy. 2008;8(11):1749–57.

81. Raveh S, Gavert N, Ben-Ze’ev A. L1 cell adhesion molecule (L1CAM) in invasive tumors. Cancer letters. 2009;282(2):137–45.

82. Roberts SA, Gordenin DA. Clustered and genome-wide transient mutagenesis in human cancers: Hypermutation without permanent mutators or loss of fitness. BioEssays: news and reviews in molecular, cellular and developmental biology. 2014;36(4):382–93.

83. Chan K, Gordenin DA. Clusters of Multiple Mutations: Incidence and Molecular Mechanisms. Annual review of genetics. 2015;49:243–67.

84. Kristjansdottir K, Fogarty EA, Grimson A. Systematic analysis of the Hmga2 3’ UTR identifies many independent regulatory sequences and a novel interaction between distal sites. RNA (New York, NY). 2015;21(7):1346–60.

