## supplemental figure 4 for ConSurf result for "Comprehensive bioinformatics analysis of *L1CAM* gene revealed Novel Pathological mutations associated with L1 syndrome"

### ConSurf Results

```
1           11          21          31          41
MVVALRYVWP  LLLCSPCLLI  QIPEEYEGHH  VMEPPVITEQ  SPRRLVVFPT
eebbbebbbb  bbbbbbbebb  ebeeeeeeee  eeeeeebeee  eeeebbbbee
f                                      ff s     f

51          61          71          81          91
DDISLKCEAS  GKPEVQFRWT  RDGVHFKPKE  ELGVTVYQSP  HSGSFTITGN
eebebebeee  eeeeeebebe  eeeeebeeee  eeebeeeeee  eeeebebebe
f     s f   fff f   s     f                      fff

101         111         121         131         141
NSNFAQRFQG  IYRCFASNKL  GTAMSHEIRL  MAEGAPKWPK  ETVKPVEVEE
eeeebeeeee  ebebebeeee  bebbeeebeb  ebeeeeeeee  eeeeeeebee
         f    fs s f    s s f         f  f   f  f

151         161         171         181         191
GESVVLPCNP  PPSAEPLRIY  WMNSKILHIK  QDERVTMGQN  GNLYFANVLT
eeebbbebee  eeeeeebbbb  bbebebeebe  eeeebeebee  bebbbbebee
f    s s f  f f         ssf     s   f  fs    f  sfs s f

201         211         221         231         241
SDNHSDYICH  AHFPGTRTII  QKEPIDLRVK  ATNSMIDRKP  RLLFPTNSSS
eeeeeebbbb  bebeebeebe  eeeebebebe  eeeeeeeeee  ebbbeeeeee
 f f fs s    fs   ffsf  ff      s            f      f    f

251         261         271         281         291
HLVALQGQPL  VLECIAEGFP  TPTIKWLRPS  GPMPADRVTY  QNHNKTLQLL
eeebbeeeeb  ebebbeeeee  eeebebeeee  eeeeeeeeee  eebeebbebe
      f       fssf f f  ff   s                      f s

301         311         321         331         341
KVGEEDDGEY  RCLAENSLGS  ARHAYYVTVE  AAPYWLHKPQ  SHLYGPGETA
ebeeeeeeeb  ebebeeeeee  bebbbebbbe  ebeebbeeee  eebbbeeeeb
     f f s   s s f  f     s   s sf  f f     f   f      f

351         361         371         381         391
RLDCQVQGRP  QPEVTWRING  IPVEELAKDQ  KYRIQRGALI  LSNVQPSDTM
ebebeeeeee  eeebebebeb  eebeeeeeee  eeebeeeebb  beebeeeeeb
   s f f f   f   s  fs                                 ffs

401         411         421         431         441
VTQCEARNRH  GLLLANAYIY  VVQLPAKILT  ADNQTYMAVQ  GSTAYLLCKA
bbebebeeee  bbbbbebbbb  bbebeeebbe  eeeeebebbe  eeebbbebeb
s fs  ff f  s    f      s    f   f       s             s

451         461         471         481         491
FGAPVPSVQW  LDEDGTTVLQ  DERFFPYANG  TLGIRDLQAN  DTGRYFCLAA
eeeeeeebeb  eeeeeeeeee  eeebebbeeb  ebebeebeee  eeeebbbebe
ffff f                          fs              f f s s

501         511         521         531         541
NDQNNVTIMA  NLKVKDATQI  TQGPRSTIEK  KGSRVTFTCQ  ASFDPSLQPS
eeeeeeebeb  ebebeeeeeb  eeeeeeeeee  eeeebebebe  eeeeeebeee
f            s     f       f                s      f

551         561         571         581         591
ITWRGDGRDL  QELGDSDKYF  IEDGRLVIHS  LDYSDQGNYS  CVASTELDVV
bebeeeeeeb  eeeeeeeebe  eeeeebebee  beeeeeeebb  bebeeebeee
  s                          s          f f     s   f  f
```

```
601
E S R A Q L L V V G
e b e b e b e b b e
              s

611
S P G P V P R L V L
e e e e e e e b e b
  f     f

621
S D L H L L T Q S Q
b e e e b b e e e e

631
V R V S W S P A E D
b e b e b b e e e e
    s   s   f f

641
H N A P I E K Y D I
e e e e b e e b b b
f f f

651
E F E D K E M A P E
e b e e e e e e e e
f     f

661
K W Y S L G K V P G
e b e e b e e e e e
              f   f

671
N Q T S T T L K L S
e e e e b e b e b e
                s

681
P Y V H Y T F R V T
e e b e b e b b b b
f       s   s s s

691
A I N K Y G P G E P
b b e e e b e e e e
    f     s   f

701
S P V S E T V V T P
e e e e e e e e e e
f     f         f

711
E A A P E K N P V D
e e e e e e e e e e
      f     f f

721
V K G E G N E T T N
b e e e e e e e e e
    f   f         f

731
M V I T W K P L R W
b e b e b e e b e e
    s   s   f

741
M D W N A P Q V Q Y
e e e e e e e b e b
      f f f       s

751
R V Q W R P Q G T R
e b e b e e e e e e
  s     f

761
G P W Q E Q I V S D
e e b e e e e b e e
    s

771
P F L V V S N T S T
b e b b b e e e e e
              f   f

781
F V P Y E I K V Q A
b b e b e b e b e b
    f           f

791
V N S Q G K G P E P
e e e e e e e e e e
  f     f     f   f

801
Q V T I G Y S G E D
e b b b b b e e e e
            f f f f

811
Y P Q A I P E L E G
e e e e b e e e b e
  f       f

821
I E I L N S S A V L
b e b b e b b b b e
        f

831
V K W R P V D L A Q
b e b e e b e e e e
    s

841
V K G H L R G Y N V
b e e e b e e b e b
    f       f

851
T Y W R E G S Q R K
b b b e e e e e e e

861
H S K R H I H K D H
e e e e e e e e e b

871
V V V P A N T T S V
b e b e e e e e e e

881
I L S G L R P Y S S
e b e e b e e b b e
        s   f   s

891
Y H L E V Q A F N G
b e b e b e b b e e
s               f

901
R G S G P A S E F T
e e e e e e e e e e
  f   f f   f

911
F S T P E G V P G H
b e e e e e e e e e
    f   f f f f

921
P E A L H L E C Q S
e e b b e b e e e e
f

931
N T S L L L R W Q P
e e e b e b e b e e
              s

941
P L S H N G V L T G
e e e e e e e b b b
f       f f

951
Y V L S Y H P L D E
b b b e b e e e e e
    s

961
G G K G Q L S F N L
e e e e e b e e e b

971
R D P E L R T H N L
e e e e e e e b e b

981
T D L S P H L R Y R
e e b e e e e e b e
                s

991
F Q L Q A T T K E G
b e b e b e e e e e
        s   f     f

1001
P G E A I V R E G G
e e e e b e e e b b
  f

1011
T M A L S G I S D F
e b e e e e e e e b
f

1021
G N I S A T A G E N
e e b b b b b e e e

1031
Y S V V S W V P K E
b b e b b b e e e e
          s

1041
G Q C N F R F H I L
e e e e b e b e b b

1051
F K A L G E E K G G
b e e e e e e e e e

1061
A S L S P Q Y V S Y
e e e e e e e e e e
                f

1071
N Q S S Y T Q W D L
e e e b b e b e e e
                  f

1081
Q P D T D Y E I H L
e e e e e b e b e b
          s

1091
F K E R M F R H Q M
b e e e e e e e e e

1101
A V K T N G T G R V
e b e e e b e e e e
      f

1111
R L P P A G F A T E
e e e e e e b b e e
                f

1121
G W F I G F V S A I
b b b b b b b b b b
  s   s       s s

1131
I L L L L V L L I L
b b b b b b b b b b
    s   s   s

1141
C F I K R S K G G K
b b b e e e e e e e
s       f f f f f f

1151
Y S V K D K E D T Q
e e b e e e e e e e
f f s f f f f     f

1161
V D S E A R P M K D
e e e e b e e e e e
  f   f   f   f f f

1171
E T F G E Y R S L E
e e e e e b e e e e
f   f   f     f

1181
S D N E E K A F G S
e e e e e e e e e e
  f   f f f

1191
S Q P S L N G D I K
e e e e e e e e e e
f f   f
```

1201 PLGSDDSLAD
eeeeeeebee
ff ff ff

1211 YGGSVDVQFN
eeeeeeeebe
f f f

1221 EDGSFIGQYS
eeeebbbeee
ffffs sf f

1231 GKKEKEAAGG
eeeeeeeeee
f

1241 NDSSGATSPI
eeeeeeeeeb
ff ff

1251 NPAVALE
ebeeeee
f f

**The conservation scale:**

| 1 | 2 | 3 | 4 | 5 | 6 | 7 | 8 | 9 |
|---|---|---|---|---|---|---|---|---|

Variable Average Conserved

e - An exposed residue according to the neural-network algorithm.
b - A buried residue according to the neural-network algorithm.
f - A predicted functional residue (highly conserved and exposed).
s - A predicted structural residue (highly conserved and buried).
X - Insufficient data - the calculation for this site was performed on less than 10% of the sequences.
